# Parental exposure to wet and dry conditions shapes the viability and thermotolerance of eggs in *Aedes aegypti*

**DOI:** 10.1101/2025.06.16.658957

**Authors:** Souvik Chakraborty, Dhruv Shah, Diya Dayal, Luke Lutz, Lyn Wang, Jazmin Garcia, Gabrielle Lefevre, Emily Susanto, Joshua B. Benoit

## Abstract

*Aedes aegypti*, a primary vector of dengue, Zika, and chikungunya, displays remarkable adaptability across ecological gradients. Central to this resilience is the egg stage, which must withstand fluctuating moisture and temperature conditions. Environmental transitions, particularly changes in moisture availability, significantly influence egg hatching success in mosquitoes. This study investigates how parental exposure to variable hydration conditions shapes key reproductive traits in *Ae. aegypti*. Using four environmental regimes, continuous wet, continuous dry, wet-to-dry, and dry-to-wet, we assessed egg output, hatching success, thermotolerance, and egg nutrient composition across three *Ae. aegypti* populations. Our results show that oviposition timing and egg production are significantly affected by the hydration environment experienced by the parental generation. While the wet, dry, and dry-to-wet groups exhibited a consistent oviposition peak beginning four days post-blood feeding, the wet-to-dry group showed delayed reproductive investment, with peak egg production occurring later. Egg output was highest under continuous wet conditions and significantly reduced in the dry and wet-to-dry treatments across all populations. Interestingly, the wet-to-dry group showed significantly higher egg-thermotolerance than any other group, and this pattern was consistent across all three populations under high-temperature stress conditions (41°C and 45°C). Nutritional composition showed an increased glycogen level in eggs when parents were exposed to wet conditions before blood feeding. By integrating physiological and ecological metrics such as hatching rates and thermal stress resilience, we demonstrate how parental environments shape subsequent egg performance, highlighting adaptive responses that enable *Ae. aegypti* persistence under increasing climate variability.

## Introduction

*Aedes aegypti* poses a significant global health threat as a vector of debilitating diseases, including dengue, Zika, chikungunya, and yellow fever. Their rapid population growth, adaptability to changing temperatures, and success across diverse ecological niches underscore their public health importance. The egg stage is central to their ecological success, with successful hatching following exposure to poor conditions as a critical factor for establishment and expansion (Benoit et al., 2023; Chu et al., 2022; Day, 2016). The ability of *Ae. aegypti* to lay viable eggs under fluctuating environmental conditions has significant implications for their persistence and dispersal (Faull et al., 2016). Fluctuations in water availability, especially cyclical wet-dry periods often driven by seasonality and climate variability, are key environmental determinants of mosquito reproduction and survival, directly affecting egg viability and hatching success (Juliano et al., 2002; Shaman and Day, 2007). Abiotic stressors, including drought or extreme humidity experienced during the parental life stages, can impact reproductive outcomes and offspring traits (Mamai et al., 2014). This kind of parental influence on offspring traits has also been observed in other arthropods, including ticks, mites, and earwigs, where some maternal effects help offspring cope with stress while others fail under extreme conditions (Le Hesran et al., 2020; Le Roux et al., 2024; Yoder et al., 2006). In particular, environmental conditions during early development stages, such as eggs, larvae, pupae, and even early adulthood, can influence offspring phenotype through altered resource allocation, developmental programming, or genetic-environmental interactions (Mousseau and Dingle, 1991). Short-term or transient environmental stress, rather than long-term exposure, can drive such responses in parents that shape offspring phenotype (Nussey et al., 2007; Whitman and Agrawal, 2009), often described as transgenerational plasticity. Emerging evidence suggests that environmental stress experienced by parents can influence reproductive physiology well before the egg is laid, reflecting a broader scope of transgenerational plasticity effect (Colloff, 1987; Sota and Mogi, 1992; Yoder et al., 2004). Many studies have investigated how extreme temperatures or humidity directly affect insect eggs, often overlooking the pre-oviposition parental environment (Fischer et al., 2003; Fox et al., 1999; Mousseau and Dingle, 1991). Parental exposure to even brief episodes of climatic stress can alter egg hatching thresholds, exemplifying transgenerational plasticity that is crucial for predicting mosquito population susceptibility and reproductive trajectories in response to erratic climatic regimes.

Maternal effect is a specific form of transgenerational plasticity that arises when the maternal genotype or phenotype shapes offspring traits in response to environmental cues, often functioning as a shared phenotype that influences both maternal and offspring fitness (J. Marshall and Uller, 2007; Rossiter, 1991; Walzer and Schausberger, 2015). Maternal effects play a crucial role in adaptive responses, allowing mothers to optimize both their survival and the performance of their offspring under environmental stress. Illustrating the impact of environmental stressors on mosquito reproductive dynamics, egg size variation in *Ae. aegypti* is directly linked to the female’s nutritional status, which dictates the allocation of crucial macronutrients, such as lipids, proteins, and glycogen, to her developing eggs (Yanchula and Alto, 2021). Effects of maternal photoperiod and diapause distinctly influence embryogenesis in the tropical and temperate strains of *Ae. albopictus (Lacour et al., 2014)*. These resources are vital during early embryonic development and can have cascading consequences for subsequent offspring fitness (Mousseau and Fox, 1998). However, resource allocation often leads to trade-offs between the number of eggs produced and their quality, ultimately shaping the evolution of life-history strategies (Mousseau and Dingle, 1991; Yanchula and Alto, 2021). While maternal effects are relatively well-studied, paternal effects are increasingly recognized, although understudied in mosquitoes. In nature, both maternal and paternal environmental conditions influence distinct aspects of offspring phenotype, including development rate, body size, and survival, sometimes in a sex-specific or conditionally dependent manner (Bonduriansky and Head, 2007; Zirbel and Alto, 2018). Paternal contributions mediated through seminal fluid proteins (sfps) or other factors can influence reproductive outcomes in the dengue vector *Ae. aegypti* (Sirot et al., 2011). Nutritionally stressed smaller males produce fewer sfps, potentially diminishing female fecundity and longevity (Helinski and Harrington, 2011). While studies on paternal larval nutrition and size have shown measurable effects, their broader influence on reproductive traits such as egg viability or stress tolerance is largely unknown. In contrast to traditional metrics such as egg size or weight, this study emphasizes egg hatching success and thermotolerance, which are ecologically relevant and quantifiable indicators of reproductive viability. By investigating how combined parental environments (pupal and adult stages) influence egg hatching performance, we offer a functional perspective on transgenerational plasticity in reproductive success. This framework enables us to evaluate how parental environmental history influences offspring resilience under heat stress, thereby enhancing our understanding of mosquito reproductive ecology in fluctuating environments.

The extent of transgenerational plasticity varies within and among populations, suggesting a potential role for genetic variation and local adaptation (Pigliucci, 2005; Sturiale and Bailey, 2023). In arthropods where water balance is crucial, such as many insects and mites, maternal effects can be significant for maintaining ecological function across diverse and often harsh environments (Gefen et al., 2006). This is true across insect taxa, where differential abiotic pressure and larval diet lead to differential resource allocation in eggs (Bonduriansky and Head, 2007; Ledón-Rettig, 2023; Macagno et al., 2018; Woestmann and Saastamoinen, 2016; Yanchula and Alto, 2021). Understanding how the parental environment influences egg viability and tolerance to environmental stressors is crucial for assessing offspring performance in variable climates. Incorporating ecologically relevant metrics such as hatching success and thermotolerance provides a more comprehensive view of parental experiences shaping egg resilience in this vector species. Moreover, parental effects, encompassing maternal and paternal influences, are not laboratory artifacts, but real, dynamic, and often reversible mechanisms by which organisms buffer against environmental unpredictability (Mousseau and Dingle, 1991). These mechanisms contribute to broader ecological outcomes such as survival, reproductive timing, and population persistence, and are becoming increasingly significant under rapidly shifting climate conditions. Transitions between environmental states, such as sudden shifts from wet to dry conditions (or vice versa), may present unique physiological challenges and adaptive opportunities.

Understanding the complex interactions between climate-associated changes and mosquito reproduction is crucial, as the global climate is reshaping mosquito habitats and influencing mosquito-human interactions worldwide. The ability of mosquito populations to thrive in these dynamic environments hinges on their capacity to adapt to environmental shifts throughout their life cycle (Benoit et al., 2023; Holmes and Benoit, 2019). While prior research has explored the effects of sustained wet or dry environments on mosquito life history, the interplay between sequential environmental experiences, particularly transitions between wet and dry conditions, remains unexplored. To address this gap, our study tests the hypothesis that emergence under wet conditions confers reproductive advantages, particularly in terms of egg output. We further examine how brief parental exposure, followed by blood feeding and a shift between wet and dry conditions, affects key reproductive traits in *Ae. aegypti*. We quantify egg number, hatching success, thermal tolerance, and egg nutrient levels across four distinct environmental regimes: continuous wet, continuous dry, wet-to-dry, and dry-to-wet. This integrative approach enables us to disentangle the effects of parental exposure to dry and wet conditions on shifts in hatching success and egg thermotolerance. Understanding how environmental transitions influence reproductive success will enhance our capacity to model mosquito population dynamics in the context of ongoing climate change.

## Materials and methods

### Study organism and experimental setup

Two populations of *Aedes aegypti* mosquitoes (PK10, Ngoye) from the West African Sahel region were used in this experiment, along with a third ‘Combined’ population created by mixing equal numbers from the two. PK10, a sparsely human-dense area with fewer than 12.5 individuals per km², contrasts sharply with Ngoye, a densely populated region exceeding 200 individuals per km², reflecting distinct environmental and climatic conditions (Chakraborty et al., 2024; Rose et al., 2020). These populations have been maintained under standard laboratory conditions of 27 ± 2°C, a relative humidity (RH) of 65–70%, and a 16:8 hour light-dark cycle since 2021. Experiments were conducted using individuals from the 14^th^ generation. Mosquitoes were routinely blood-fed on human hosts (University of Cincinnati IRB 2021-0971), and oviposition cups lined with brown, hardwound paper were placed inside mosquito cages. These papers were collected two weeks after blood feeding and stored wrapped in moist paper towels inside sealed ziplock bags under laboratory conditions. Two weeks after egg harvesting and storage, the eggs were placed in pans (30.5 cm × 7.5 cm × 5 cm) containing a rearing medium of deionized water, finely ground Tetramin Goldfish Flakes, and yeast extract. Approximately 200 larvae were reared per pan to maintain consistent density. Upon pupation, pupae were transferred with a pipette into cups and subsequently placed in mesh cages (30 cm×30 cm×30 cm), marking day 0 (**Figure 1**). The cages were housed under either “wet” or “dry” conditions. The wet condition corresponded to the climate-controlled laboratory environment (mentioned previously), and the dry condition was maintained in a separate room set to 33 ± 2 °C, 33–38% RH, and the same light-dark cycle. In all experimental cages, test tubes containing cotton wicks soaked in water and 10% sucrose solution were provided for 8 hours during the light phase starting from day 1, as adult emergence. All populations received the same access to sugar water unless otherwise specified. On day 8, mosquitoes were blood-fed under wet and dry conditions, with 10 females per replicate (n = 10) per population. An additional set of mosquitoes was also blood fed, and following blood feeding, a reciprocal transfer ("vice versa") of wet and dry conditions was performed, where subsets of mosquitoes originally reared in dry or wet environments were moved to the opposite condition (**Figure 1**). This allows us to evaluate the impact of four different conditions, i.e., (continuous) wet, (continuous) dry, wet-(transitioning)- to-dry, and dry-(transitioning)-to-wet, on five replicates of ten mosquitoes each. Survival and reproductive output were monitored through periodic sampling on days 12, 16, and 20 to assess survival progression and oviposition egg numbers.

**Figure 1:**
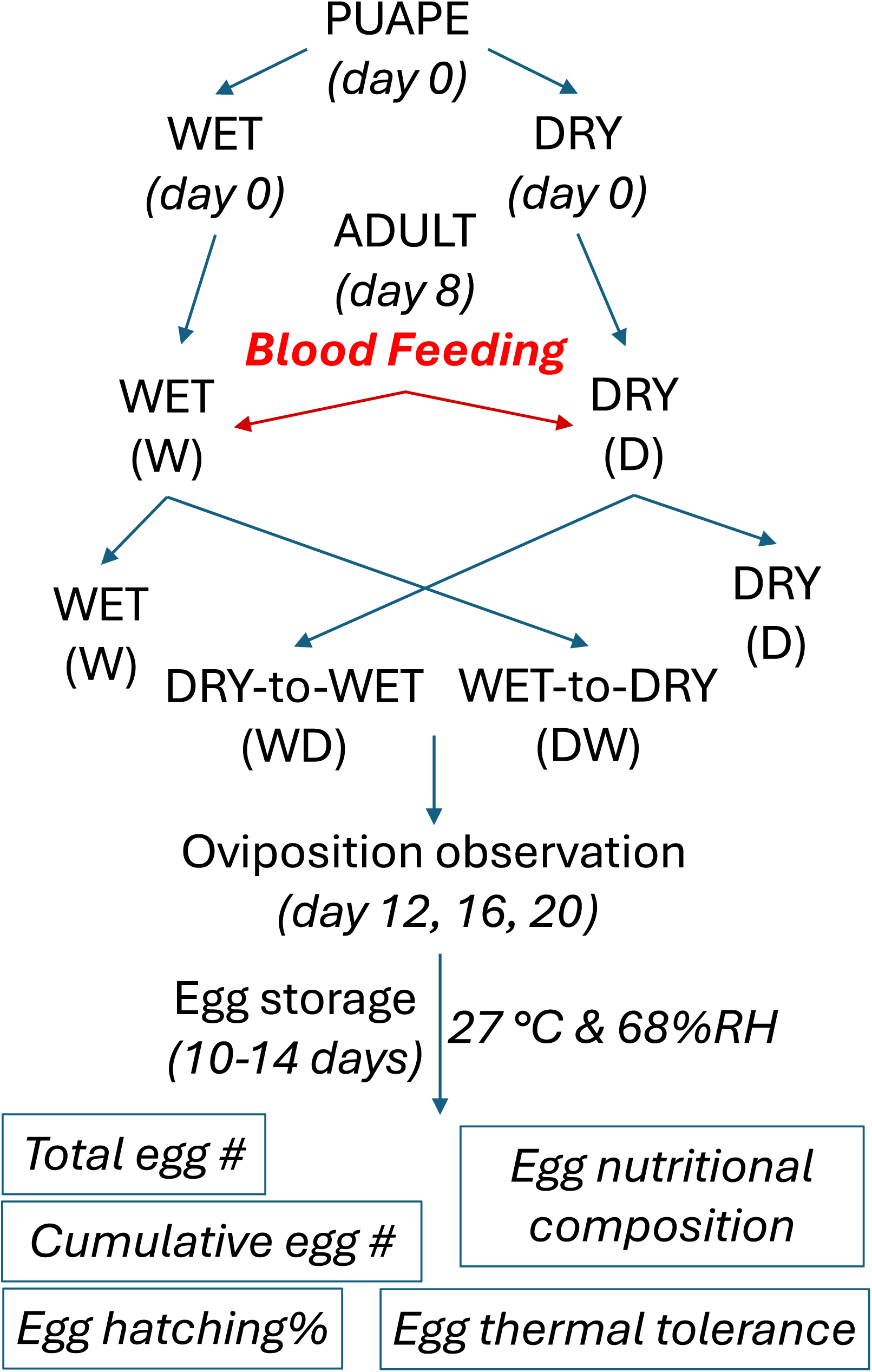
Summary of experimental procedure. Pupae from three mosquito populations (PK10, Ngoye, and Combined) were placed in cages maintained under either wet (27 ± 2 °C, 65–70% RH) or dry (33 ± 2 °C, 33–38% RH) conditions. Eight days after blood-feeding, females were subjected to a reciprocal transfer, resulting in four distinct hydration regimes. Eggs were collected at 4-day intervals post-blood-feeding (days 12, 16, and 20). The total number of eggs, egg hatching percentage, egg thermotolerance, and egg nutritional composition were assessed. Each treatment group included five biological replicates, with 10 females per replicate.

### Transgenerational effect quantification matrix

We calculated four indices to assess the trade-off associated with parental exposure to different levels of wet and dry conditions: (i) temporal progression of oviposition, (ii) egg hatching success, (iii) egg thermotolerance, and (iv) egg nutritional composition.

(i) The hardwound paper from the oviposition cups was collected, and egg numbers were counted every four days following blood-feeding on day 8 (age 12, 16, and 20). Adult female survival rates were recorded, and the average egg progression was plotted as oviposition per mosquito. Two weeks following storing the eggs, egg hatching success and egg thermotolerance were examined, where batches of 30 - 70 eggs were placed in 8 mL plastic vials (Thornton Plastics: 2.5 Udbl), with 12 replicates per treatment. (ii) A set of replicates filled with DI water and maintained under standard laboratory conditions (29 °C) was used to assess egg hatching success under normal conditions.

(iii) Thermal stress was applied to the eggs at 33 ± 1, 37 ± 1, 41 ± 1, and 45 ± 1 °C for 4 hours. The vials (12 replicates per treatment) were placed in dry heat blocks inside a digital dry bath (Thermo Scientific™) and covered with styrofoam to maintain a constant temperature throughout the experiment. A 30-minute temperature ramp preceded the stress, starting the heat bath at 29 °C. During the initial 15 minutes, the temperature was adjusted to the midpoint between 29 °C and the target temperature, and during the subsequent 15 minutes, it was further adjusted to reach the target temperature. After the thermal stress treatment, the egg vials were filled with DI water and fish food once the temperature returned to 29 °C. Forty to forty-eight hours after the experiment, the eggs and larvae were preserved with 80% ethanol, and the vials were stored at -20 °C until viability counts were conducted. Proportional survival rates were compared to evaluate the effects of thermal stress on egg thermotolerance. To quantify the transgenerational impact on reproductive success following parental exposure to varying wet-dry conditions, we developed a composite metric defining Transgenerational Efficacy Index (TEI), integrating two key parameters, i.e., the average number of eggs laid per surviving female (fecundity), and mean proportion of eggs hatched following exposure to elevated thermal stress at 41 °C and 45 °C (egg thermotolerance). The TEI was calculated as the product of these two values, effectively combining maternal investment and embryonic resilience into a single, interpretable measure of reproductive performance under abiotic stress, with potential applicability across other ectothermic organisms.

(iv) Two weeks post-storing the egg set of eggs, nutrient quantification assays for lipids, glycogen, and proteins were performed using lab-standardized protocols (Bailey et al., 2024; Holmes et al., 2023) on the Combined population only, as prior analyses showed no significant differences in adult female reserves or reproductive indices across populations. Consistency was maintained across three technical and three biological replicates. Ten days after storing eggs under ambient conditions (29 °C), a subset of eggs were flash-frozen at –80 °C for 30 minutes. Groups of fifty eggs (n = 50) were then homogenized in STE buffer containing 2% Na_2_SO_4_ using a BeadBlaster 24 (Benchmark), and the homogenate was divided into aliquots for lipid (100 μL), trehalose (150 μL), and glycogen (150 μL) analyses. Assays were run across two 96-well plates (Zinsser), with each biological group represented in triplicate. Two standard curves per plate, each in technical duplicate, were included for calibration. Lipid levels were quantified by absorbance at 525 nm, glycogen levels were measured at 625 nm, and protein levels were measured at 595 nm using a Synergy H1 microplate reader (BioTek). An additional set of eggs (n = 20) was placed in a desiccator (Fisher Scientific 100L Oven Grvty) for two weeks to determine dry mass. Standard curves for all macronutrient measurements were normalized to average egg mass and individual egg count to account for weight differences between samples.

### Statistical analyses

Linear regression models using the ’lm’ function in the lme4 package were applied to analyze differences in egg production, viability, and nutritional reserve across experiments. Post-hoc comparisons were performed with the "emmeans" package, and statistical significance was determined at *p* < 0.05. Statistical analyses were conducted in RStudio (version 4.3.2) and figures were edited in Inkscape (version 1.3.2).

## Results

### Temporal progression of oviposition

Egg production varied by hydration condition and time post-blood feeding across populations (**Figure 2**). In the wet, dry, and dry-to-wet conditions, oviposition generally followed a unimodal pattern, with peak egg output occurring between 4-8 days post-blood feeding (day 12- 16). While cumulative egg output differed by population across these treatments, the overall temporal pattern remained consistent. In contrast, the wet-to-dry group showed a markedly different trajectory. Egg production remained low during the first two intervals (0-4 and 4-8 days), with a delayed and pronounced peak emerging only between 8-12 days post-blood feeding (age 16-20), suggesting a lag in reproductive investment under transitional stress. Interestingly, during the 4-8 day period post-blood-feeding, the Ngoye population produced significantly more eggs under the wet-to-dry condition compared to other PK10 and ‘Combined’ (F_2,12_ = 7.96, *p* = 0.006; **Figure 2a**). Except for this subtle case, no influence of population on egg production was apparent for any of the treatment conditions. Overall, only treatment condition exerted a strong, highly significant influence on egg production (F_3,176_ = 13.25, *p* = 7.69྾10^-8^).

**Figure 2:**
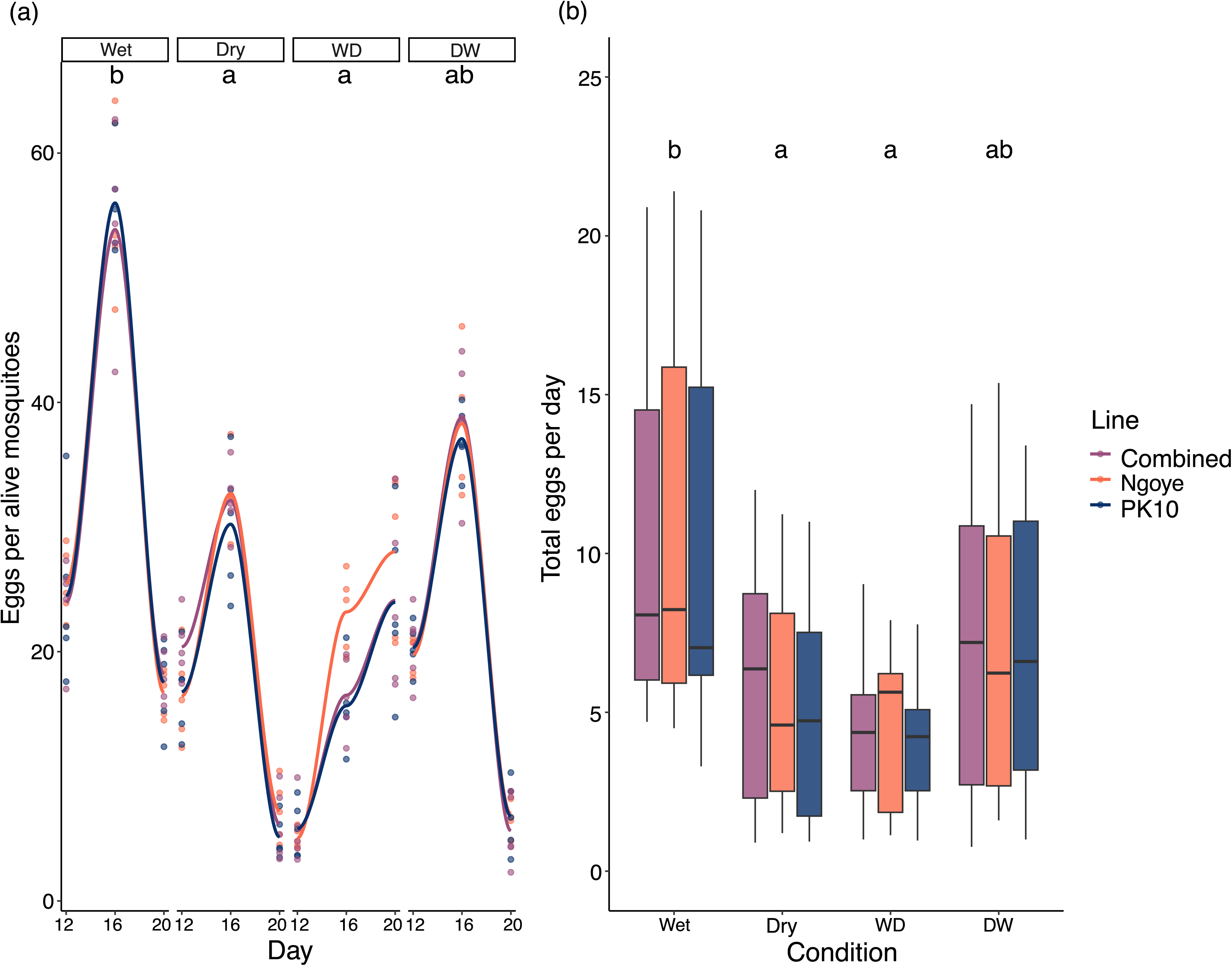
Temporal dynamics of egg production across hydration treatments. (a) Eggs laid per surviving female recorded at a four-day interval following blood feeding, and (b) total egg production per day by a surviving female across four hydration conditions. Each trial was initiated with five biological replicates (n = 5), each containing 10 females. Survival counts were recorded every four days, and daily egg production was normalized by the number of surviving females. Letters indicate statistically significant differences between treatments (*p* < 0.05), as determined by ANOVA followed by pairwise comparisons using post-hoc tests.

Relative to the wet condition, reduced egg production under constant dry and wet-to-dry conditions was consistent for all three populations studied (**Supplementary Table 1**). Daily egg production per female followed the temporal progression (**Figure 2b**), revealing distinct differences in egg-laying patterns across conditions.

### Egg hatching responses to varying levels of wet and dry conditions

Egg hatching success varied significantly across hydration treatments (F_3,140_ = 5.872, *p* = 0.0008), although hatching rates consistently exceeded the 80% threshold. Overall, mean hatching success was highest under continuously wet conditions and declined in the dry, wet-to-dry, and dry-to-wet groups. Tukey post hoc comparisons revealed that the strongest reductions occurred under dry (*p* = 0.0035) and dry-to-wet (*p* = 0.0007) conditions relative to the wet treatment. In contrast, no significant difference was observed between the wet and wet-to-dry treatments (*p* = 0.06). However, hatching under dry conditions did not differ significantly from either transitional treatments (dry-to-wet or wet-to-dry), indicating a general decline in viability under any deviation from full hydration (**Figure 3**). There was minimal significance of population effect on egg hatching (F_2, 132_ = 4.662, *p* = 0.01106), and the interaction between population and hydration condition on egg hatching was not significant (_F6, 132_ = 0.691, *p* = 0.65686) (**Supplementary Table 2)**. Across all conditions, no significant population-level differences were detected. However, under the wet-to-dry condition, the ‘Combined’ population exhibited a higher hatching rate than both founder populations (PK10 and Ngoye) (**Figure 3**).

**Figure 3:**
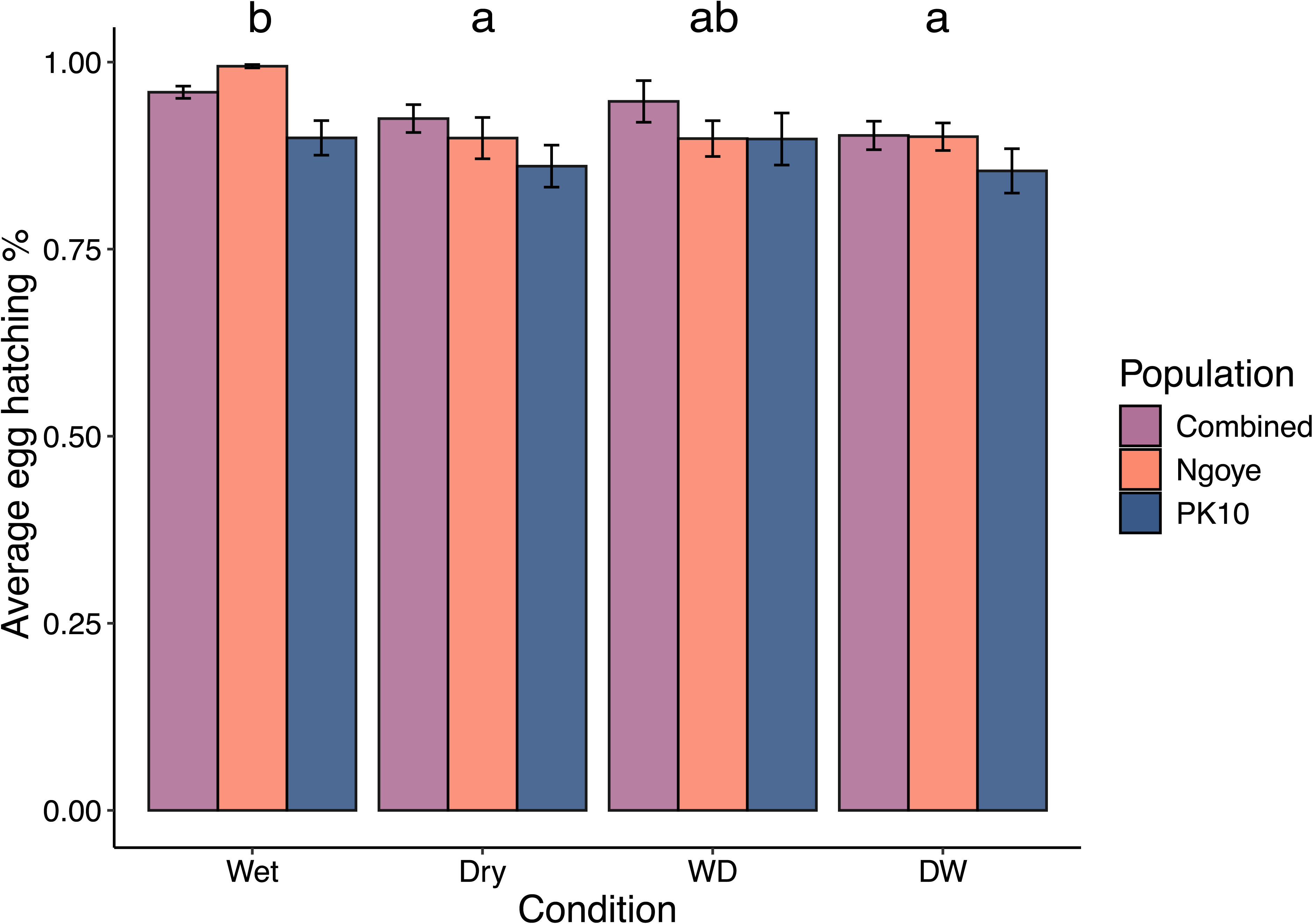
Egg hatching success across hydration treatments. Proportion of eggs hatched under standard laboratory conditions (29 °C, 65–70% RH) from parents subjected to different hydration treatments. Each treatment group included 12 replicates, with 30– 70 eggs per replicate. Letters indicate statistically significant differences in hatching success between treatments (*p* < 0.05), based on ANOVA followed by post-hoc comparisons (emmeans). Proportional hatching success was calculated as Hatched eggs / (Hatched eggs + Unhatched eggs).

### Variation in egg thermotolerance among strains exposed at varying levels of wet and dry conditions

Egg thermotolerance varied significantly across hydration treatments (F_3,572_ = 4.19, *p* = 0.006) at all tested temperatures, with treatment effects becoming increasingly pronounced at higher temperatures. Differences among hydration treatments were detectable even under mild thermal stress (33 °C and 37 °C), but these disparities became increasingly marked as temperatures rose, underscoring the compounding impact of extreme heat on egg hatching success (**Supplementary Table 3**). While egg thermotolerance varied across populations and conditions, a consistent hatching rate above 70% at 33°C and 37°C sharply declined to below 25% at 41°C and 45°C, with only a few exceptions (**Figure 4**). The "Combined" population experiencing wet-to-dry transitional conditions exhibited 0.31 ± 0.09% and 0.16 ± 0.01% of egg thermotolerance under 41°C and 45°C, respectively **(Supplementary Table 4)**. Notably, under these high temperatures, eggs from this particular transitional condition consistently showed significantly greater thermotolerance than those from dry-to-wet, dry, and even wet groups, suggesting a robust protective effect of rehydration (**Figure 4**). At 45 °C, the wet-to-dry group outperformed all others, including the Wet group, indicating that exposure to wet conditions before exposure to dryness may confer enhanced resilience to heat exposure. When population and hydration effects were analyzed together, only the hydration condition remained a significant predictor of thermotolerance (F_3,564_ = 4.16, *p* = 0.006). Neither population nor the interaction term showed significance, suggesting largely treatment-driven and population-independent effects.

**Figure 4:**
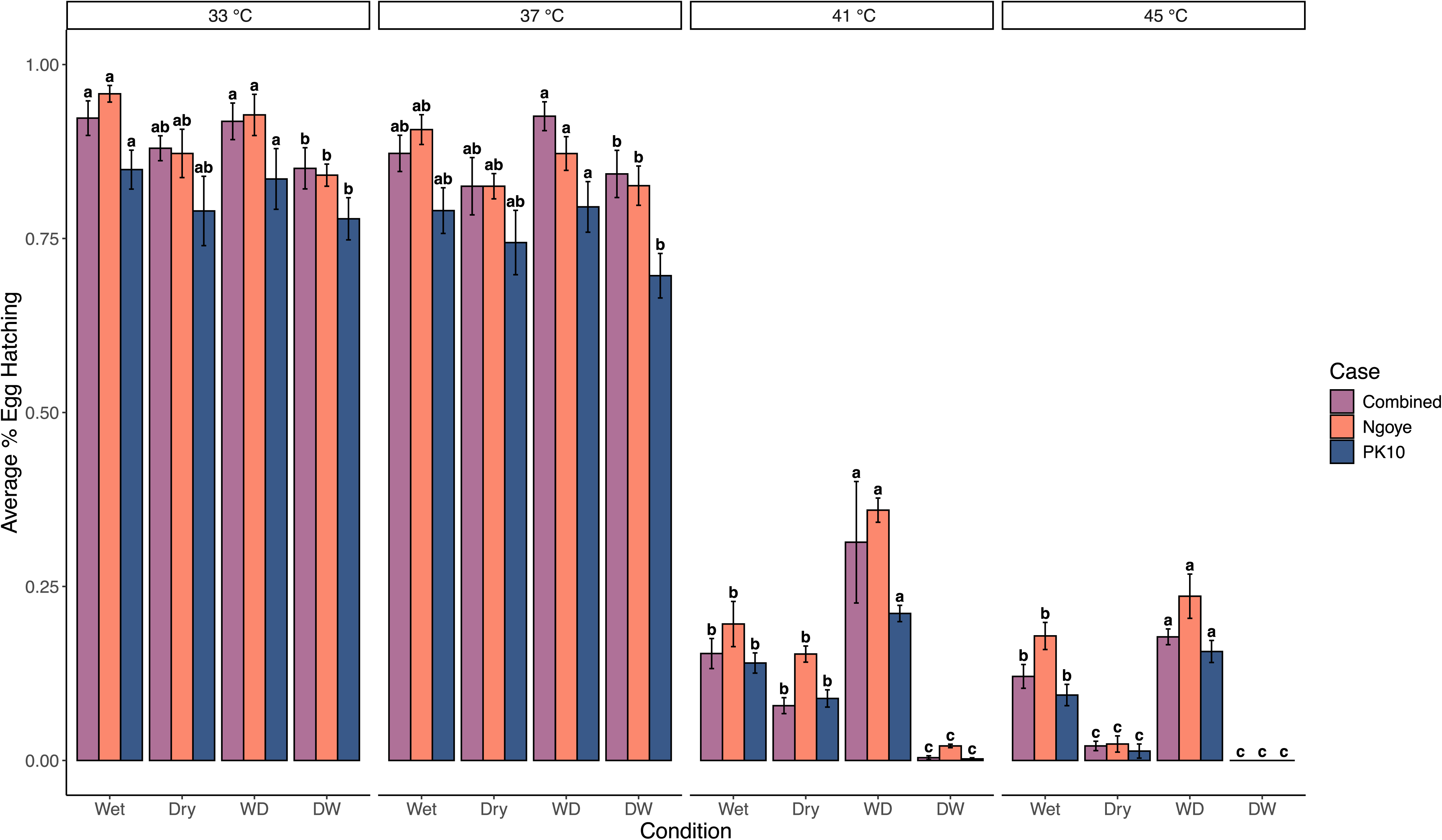
Egg thermotolerance across hydration treatments under increasing thermal stress. Eggs from parents exposed to each of the four hydration treatments were subjected to 4-hour heat stress at 33 °C, 37 °C, 41 °C, and 45 °C. Each temperature–treatment combination included 12 replicates, with 30–70 eggs per replicate. Box plots with standard error bars show the distribution of proportional hatching success. Statistically significant differences in hatching success between treatments were identified at *p* < 0.05, based on ANOVA followed by post-hoc comparisons (emmeans).

To further investigate transgenerational responses under thermal stress, we calculated a Transgenerational Efficacy Index (TEI), integrating both fecundity (average eggs laid per surviving female) and offspring viability (egg thermotolerance at 41°C and 45°C) (**Figure 5**). Across all populations, TEI patterns were consistent, where the highest efficacy was observed under constant wet conditions, with Ngoye showing the highest TEI overall. A marked decline in TEI was observed under dry and dry-to-wet treatments, indicating strong suppression of transgenerational performance under persistent or earlier dehydration stress in the life cycle. The wet-to-dry condition produced intermediate TEI values, suggesting partial buffering of maternal effects when initial hydration was favorable. For nutrient levels, no statistically significant differences were observed in egg dry mass or macronutrient content across conditions, except for a single significant effect in glycogen levels between the dry and wet-to-dry treatments (*p* = 0.03) (**Supplementary Figure 1**).

**Figure 5:**
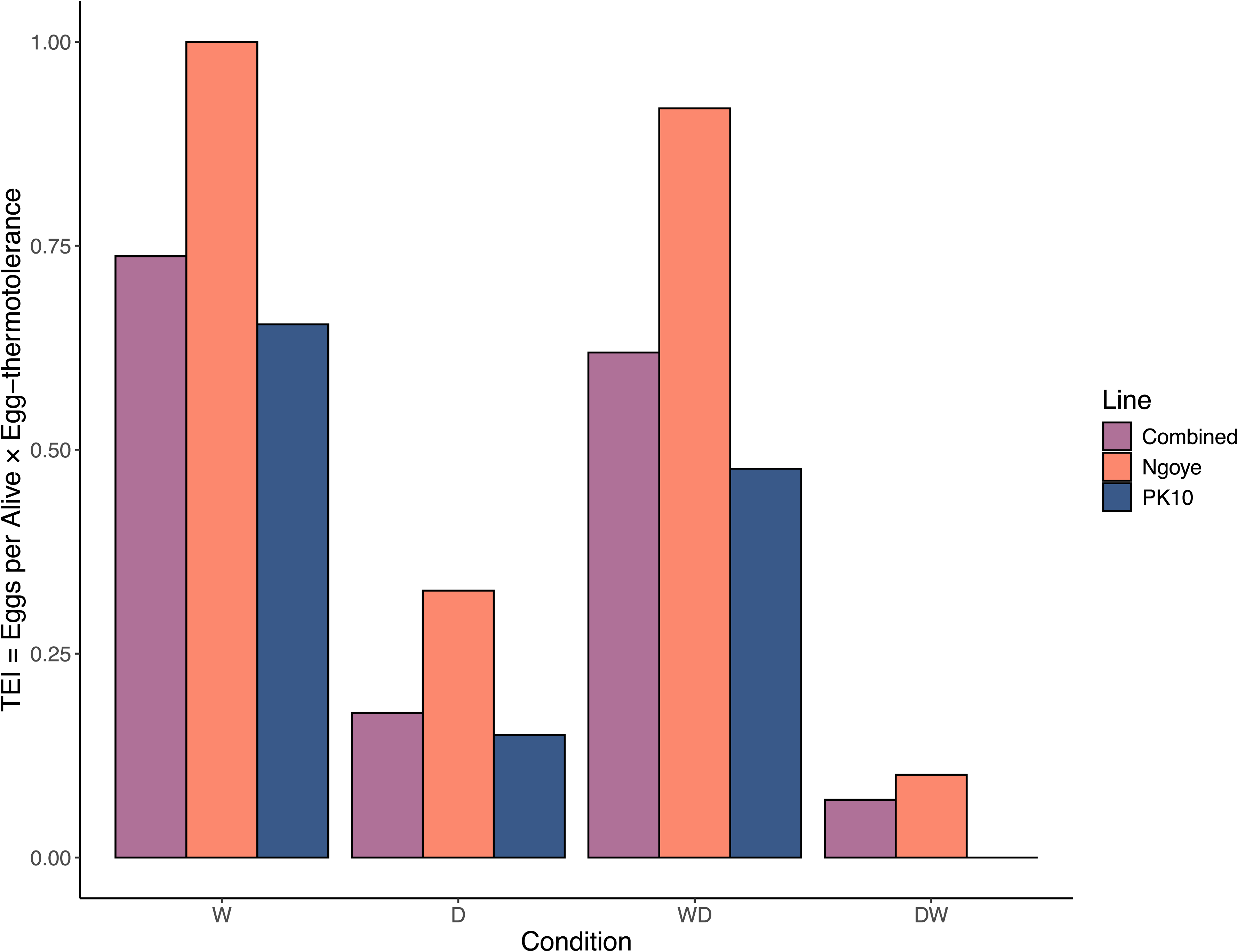
Transgenerational efficacy index (TEI) outlining fecundity and thermotolerance. A composite metric of transgenerational efficacy, calculated as the product of egg production and thermotolerance, across hydration treatments for *Aedes aegypti* females from three populations. TEI values reflect offspring’s combined reproductive output and stress resilience under varying maternal environments.

## Discussion

Understanding how environmental variability influences reproductive traits is crucial for predicting the population trajectories of vector species, such as *Aedes aegypti,* under ongoing climatic shifts. While sex-specific and context-dependent parental effects on life history traits are well-documented in insects (Bonduriansky and Head, 2007; Dew-Budd et al., 2016; Zanchi et al., 2011), the consequences of fluctuating wet and dry environments on mosquito reproduction remain understudied. The transient and unpredictable nature of mosquito breeding sites, often ephemeral aquatic habitats, forces them to navigate complex hydrological challenges that influence both survival and reproductive strategies (Day, 2016). In this study, we evaluated the cyclic impacts of wet and dry conditions on key reproductive traits. Parental exposure to a wet-to-dry transition resulted in the lowest egg output; however, these eggs exhibited enhanced thermotolerance, suggesting a potential adaptive response to increasing environmental dryness.

Hydration conditions profoundly shaped the temporal trajectory of egg production in *Ae. aegypti*, revealing plasticity in reproductive timing in response to environmental stress. While constant wet, dry, and dry-to-wet environments supported a predictable, unimodal oviposition peak at 4-8 days post-blood feeding, exposure to wet-to-dry conditions resulted in a delayed reproductive event. This temporal lag aligns with findings that females often delay reproduction following exposure to variable environments, optimizing egg-laying strategies to maximize fitness under fluctuating conditions (Hilker et al., 2023; Le Hesran et al., 2019). This shift may be an adaptive maternal strategy, as oviposition is entirely controlled by the mother, allowing her to delay reproduction until more favorable environmental cues are present, thereby enhancing offspring viability (Benton et al., 2005; Mousseau and Dingle, 1991). Short-term environmental mismatches can significantly impact reproductive decisions, particularly in species with limited mobility during early life stages (Potter and Woods, 2012; Schausberger, 1998). In mosquitoes, postponed egg-laying may represent a form of reproductive risk-spreading or bet-hedging, consistent with the broader concept of maternal effects, where females adjust their reproductive strategies to retain eggs longer, preventing their exposure to adverse conditions (Edgerly et al., 1998; Hopper, 1999). Retaining eggs longer can be an adaptive maternal response to fluctuating cues, where females prioritize survival over fecundity by retaining eggs, aided by ovary-specific genes like tweedledee and tweedledum. Prolonged egg retention, driven by ovary-specific genes like tweedledee and tweedledum, appears to be an adaptive maternal strategy in response to fluctuating environmental cues, enabling females to prioritize survival over fecundity (Venkataraman et al., 2023). Similarly, under desiccating conditions, females often engage in secondary blood feeding to maintain hydration, highlighting a shift toward survival and delayed reproduction over immediate egg output (Holmes et al., 2025). Interestingly, the urban-origin population, Ngoye, exhibited a more pronounced late-phase oviposition response under wet-to-dry conditions, suggesting local adaptation or population-level differences. Although the influence of urbanization was not a prominent or specifically evaluated factor, these findings highlight the importance of considering not reproductive output timing, a critical yet often overlooked aspect of fitness that is shaped by parental experience in variable environments.

Hydration conditions influenced not only egg production but also had a marked effect on hatching success in *Ae. aegypti.* As expected, eggs laid under consistently wet conditions exhibited the highest hatching rates. In contrast, hatching success declined markedly when adults were exposed to continuous dry or dry-to-wet transitional environments. Prior research has established that high-moisture substrates promote optimal oviposition and embryonic development (Rezende et al., 2008; Saifur et al., 2010; Sota and Mogi, 1992), while high temperatures can facilitate embryogenesis, provided that minimal moisture is maintained to support gas exchange and metabolic activity (Bethke and Redak, 1996). Increasing temperatures have been linked to accelerated oviposition and increased egg deposition, with some species exhibiting higher egg loads at elevated temperatures (Hans et al., 2019). Notably, eggs from the wet-to-dry group exhibited hatching success nearly comparable to the fully wet group, suggesting that favorable initial conditions can partially buffer against later desiccation. Our findings underscore that *Ae. aegypti* reproduction is sensitive not only to absolute moisture levels but also to the stability and predictability of environmental conditions (Kearney et al., 2009; Mogi and Tuno, 2014; Reinhold et al., 2018). This is further supported by evidence that even minor fluctuations in temperature and humidity can profoundly influence oviposition timing, fecundity, and survival in *Ae. aegypti,* with higher temperatures coupled with lower humidity, significantly reduce reproductive success and egg viability (Costa et al., 2010). While some degree of survival under fluctuating moisture may be sufficient to sustain low-level transmission, this tolerance could enable persistent vector populations and elevate outbreak potential in the face of increasingly variable climate scenarios. We found no significant interaction between population and hydration treatment for hatching success at the control temperature. This consistency suggests that, in contrast to the subtle population-level differences in the temporal progression of egg output, sensitivity to hydration stress is likely a conserved trait across *Ae. aegypti* populations. Comparable maternal effects on egg drought resistance have been documented across arthropods, suggesting a conserved physiological response (Le Hesran et al., 2020, 2019; Yoder et al., 2006). Studies under laboratory selection show that desiccation tolerance can evolve rapidly, with consistent patterns observed across strains despite genetic variation (Hoffmann et al., 2001; Telonis-Scott et al., 2006).

Beyond fecundity and hatching rates, our study examined whether *Ae. aegypti* parental hydration status influences the thermotolerance of eggs, a trait of increasing relevance under global climate change scenarios (Chakraborty et al., 2024; Zhou et al., 2018). We found that prior adult exposure to hydration stress significantly shaped egg resilience to thermal extremes, with notable inter-treatment differences in survival at elevated temperatures (41 °C and 45 °C). Eggs derived from adults under dry or dry-to-wet conditions were generally more vulnerable to heat stress, suggesting paternal exposure to dry conditions may impair subsequent egg thermotolerance. These findings highlight that hydric and thermal stress will both impact female physiology and oviposited eggs, suggesting dehydration may impair physiological mechanisms required for heat tolerance, such as membrane stability, enzyme function, or water conservation (Benoit et al., 2010b; Farnesi et al., 2017). Most likely, processes such as membrane lipid remodeling (Prasad et al., 2023), cuticular hydrocarbon difference (Drijfhout et al., 2009), eggshell melanization (Farnesi et al., 2017), and serosal cuticle formation (Vargas et al., 2014) are likely altered by dry conditions that can lower thermal tolerance. A striking result emerged in the wet-to-dry group, despite experiencing drought in the later stages of adult life; their eggs displayed the highest thermotolerance, even exceeding that of consistently hydrated controls. This pattern may suggest a form of stress hardening, whereby an initial buffered wet phase profoundly primes the pharate larvae for subsequent environmental extremes (Benoit et al., 2010a; Li et al., 2009; Lopez-Martinez et al., 2009; Sinclair et al., 2007). This effect appeared to be explicitly triggered in groups where females experienced dryness after blood feeding.

The superior thermotolerance of the wet-to-dry groups suggests that rehydration followed by dehydration may serve as a physiological cue for maternal provisioning, thereby enhancing embryo resilience. This aligns with broader maternal effects frameworks, in which parents modulate offspring traits based on environmental predictability to enhance fitness (Burgess and Marshall, 2014; J. Marshall and Uller, 2007). Although both parents experienced hydration stress, the timing and proximity of maternal desiccation to oviposition, especially compared to the earlier exposure, likely play a dominant role in shaping egg thermotolerance, underscoring the critical influence of maternal conditioning. These findings highlight that the sequence of environmental stressors can shape offspring performance in ways that promote resilience (Hilker et al., 2023), an insight critical for predictive modeling of population outbreaks. Our results support the idea that exposure to drought following blood feeding may activate physiological priming pathways in *Ae. aegypti* females, consistent with the combined roles of within-generation and transgenerational plasticity, enhancing offspring stress tolerance under unpredictable environmental conditions (Sturiale and Bailey, 2023). Notably, the high variability in egg hatching observed under wet-to-dry conditions (∼12% to 80%) for the Combined population may reflect a strategy wherein maternal exposure to fluctuating environments produces a heterogeneous set of offspring phenotypes. Such variability could arise from plastic responses that diversify thermal or dehydration tolerance among eggs, potentially increasing survival across spatially or temporally unpredictable microhabitats. This pattern parallels findings from insect taxa inhabiting ephemeral habitats, such as *Aedes triseriatus*, where females respond to seasonal cues by altering their oviposition behavior, laying eggs more opportunistically when offspring are likely to enter diapause later in the season (Edgerly et al., 1998). Reduced immediate reproductive output coupled with increased offspring stress resilience may reflect a fitness-maximizing tradeoff under uncertain conditions (Crean and Marshall, 2009; Simons, 2011). During dry spells, temporary larval pools often warm up, increasing the likelihood of heat and dehydration risks occurring simultaneously. In these situations, laying fewer but stress-tolerant eggs could improve population survival in erratic microhabitats, revealing maternal decisions to optimize future offspring survival in response to environmental unpredictability, conceptually aligning with bet-hedging strategies (Hopper, 1999).

The observed depletion in glycogen in eggs laid by continuously dry-reared mosquitoes reflects a maternal stress response, likely reducing investment in carbohydrate provisioning under conditions of desiccation. This mirrors the patterns observed in western flower thrips, *Frankliniella occidentalis*, where adult dehydration resulted in substantial depletion of glycogen reserves, particularly in females, and was associated with increased expression of genes involved in carbohydrate metabolism (Bailey et al., 2024). Notably, glycogen levels were highest in the wet-to-dry group, suggesting that mothers reared under benign conditions accumulated sufficient energy reserves before encountering dehydration stress during the egg-laying period. Conversely, the dry-to-wet group exhibited intermediate glycogen levels, indicating partial recovery of maternal energy status after relief from dehydration stress following blood feeding. These contrasting patterns highlight the importance of stress timing in relation to maternal provisioning and may indicate a dynamic, condition-dependent trade-off between maternal survival and offspring investment. While no widespread changes were observed in protein or lipid content across treatments, the selective effect on glycogen may indicate an adaptive response prioritizing metabolic water generation during desiccation stress (Benoit et al., 2005; Loveridge and Bursell, 1975; Rosendale et al., 2016). These subtle but specific alterations in maternal provisioning may contribute to transgenerational adjustments in egg drought tolerance, highlighting the need for further research into the molecular and developmental consequences of parental hydration status.

In tropical and subtropical regions, where environmental conditions oscillate unpredictably between wet and dry periods, the ability of *Ae. aegypti* to adjust reproductive timing, produce stress-hardened eggs, and sustain hatching success may be critical for population persistence. Our findings suggest that *Ae. aegypti’*s reproductive strategy is not solely shaped by the magnitude of environmental stress but also by its temporal dynamics, as highlighted in the wet-to-dry group eggs. This highlights the importance of accounting for temporal variability in stress exposure when evaluating mosquito life history and resilience, a pattern also observed in other arthropods (Potter and Woods, 2012; Schausberger, 1998), where fine-scale environmental changes contribute to local adaptation. Understanding how *Ae. aegypti* eggs respond to dynamic and fluctuating climatic conditions can enhance predictions of population stability under future climate scenarios. From a public health perspective, the ability of eggs to withstand and hatch after environmental stress, such as drought followed by rainfall, may lead to sudden population surges, complicating mosquito control efforts.

## Acknowledgements

This study was partially supported by the National Institute of Allergy and Infectious Diseases of the National Institutes of Health under Award Number R01AI148551 and R21AI166633.

---

**Supplementary Figure 1: Egg macronutrient levels across hydration treatments.**

Relative glycogen (a), protein (b), and lipid (c) content in eggs laid by mosquitoes exposed to different hydration conditions. Each bar represents the average macronutrient content per 50-egg sample across three biological and three technical replicates. Nutrient levels were quantified using standardized colorimetric assays and normalized to average egg mass. Statistical comparisons were conducted using one-way ANOVA with post-hoc Tukey tests.

**Supplementary Table 1. Tukey’s post hoc comparisons of cumulative egg production across hydration treatments.**

Following ANOVA indicating a significant effect of hydration condition (but not population) on egg laying, pairwise contrasts show differences in egg production across treatment conditions.

**Supplementary Table 2. Tukey’s post hoc comparisons of egg hatching across populations following hydration treatments.**

Following ANOVA indicating a minor population effect and no interaction between population and hydration condition, pairwise contrasts assess differences in egg hatching success from parents exposed to different hydration conditions.

**Supplementary Table 3. Tukey’s post hoc comparisons of egg thermotolerance across hydration treatments at each tested temperature.**

Pairwise contrasts reveal differences in egg thermotolerance among hydration conditions across four heat stress levels (33 °C, 37 °C, 41 °C, 45 °C).

**Supplementary Table 4. Average egg thermotolerance across populations, hydration conditions, and temperatures.**

Mean ± SEM egg hatching success across three populations, four hydration conditions, and four test temperatures.

## References

1. Bailey, S.T., Kondragunta, A., Choi, H.A., Han, J., Rotenberg, D., Ullman, D.E., Benoit, J.B., 2024. Dehydration yields distinct transcriptional shifts associated with glycogen metabolism and increases feeding in the western flower thrips, *Frankliniella occidentalis*. Entomol. Exp. Appl. 172, 154–167.

2. Benoit, J.B., Lopez-Martinez, G., Phillips, Z.P., Patrick, K.R., Denlinger, D.L., 2010a. Heat shock proteins contribute to mosquito dehydration tolerance. J. Insect Physiol. 56, 151–156.

3. Benoit, J.B., McCluney, K.E., DeGennaro, M.J., Dow, J.A.T., 2023. Dehydration Dynamics in Terrestrial Arthropods: From Water Sensing to Trophic Interactions. Annu. Rev. Entomol. 68, 129–149.

4. Benoit, J.B., Patrick, K.R., Desai, K., Hardesty, J.J., Krause, T.B., Denlinger, D.L., 2010b. Repeated bouts of dehydration deplete nutrient reserves and reduce egg production in the mosquito *Culex pipiens*. J. Exp. Biol. 213, 2763–2769.

5. Benoit, J.B., Yoder, J.A., Rellinger, E.J., Ark, J.T., Keeney, G.D., 2005. Prolonged maintenance of water balance by adult females of the American spider beetle, *Mezium affine* Boieldieu, in the absence of food and water resources. J. Insect Physiol. 51, 565–573.

6. Benton, T.G., Plaistow, S.J., Beckerman, A.P., Lapsley, C.T., Littlejohns, S., 2005. Changes in maternal investment in eggs can affect population dynamics. Proc. Biol. Sci. 272, 1351– 1356.

7. Bethke, J.A., Redak, R.A., 1996. Temperature and moisture effects on the success of egg hatch in *Trirhabda geminata* (Coleoptera: Chrysomelidae). Ann. Entomol. Soc. Am. 89, 661–666.

8. Bonduriansky, R., Head, M., 2007. Maternal and paternal condition effects on offspring phenotype in *Telostylinus angusticollis* (Diptera: Neriidae). J. Evol. Biol. 20, 2379–2388.

9. Burgess, S.C., Marshall, D.J., 2014. Adaptive parental effects: the importance of estimating environmental predictability and offspring fitness appropriately. Oikos 123, 769–776.

10. Chakraborty, S., Zigmond, E., Shah, S., Sylla, M., Akorli, J., Otoo, S., Rose, N.H., McBride, C.S., Armbruster, P.A., Benoit, J.B., 2024. Thermal tolerance of mosquito eggs is associated with urban adaptation and human interactions. bioRxivorg. 10.1101/2024.03.22.586322

11. Chu, E., Chakraborty, S., Benoit, J.B., DeGennaro, M., 2022. Chapter 25: Water homeostasis and hygrosensation in mosquitoes, in: Sensory Ecology of Disease Vectors. Brill | Wageningen Academic, The Netherlands, pp. 655–682.

12. Colloff, M.J., 1987. Differences in development time, mortality and water loss between eggs from laboratory and wild populations of Dermatophagoides pteronyssinus (Trouessart, 1897) (Acari: Pyroglyphidae). Exp. Appl. Acarol. 3, 191–200.

13. Costa, E.A.P. de A., Santos, E.M. de M., Correia, J.C., Albuquerque, C.M.R. de, 2010. Impact of small variations in temperature and humidity on the reproductive activity and survival of *Aedes aegypti* (Diptera, Culicidae). Rev. Bras. entomol. 54, 488–493.

14. Crean, A.J., Marshall, D.J., 2009. Coping with environmental uncertainty: dynamic bet hedging as a maternal effect. Philos. Trans. R. Soc. Lond. B Biol. Sci. 364, 1087–1096.

15. Day, J.F., 2016. Mosquito oviposition behavior and vector control. Insects 7, 65.

16. Dew-Budd, K., Jarnigan, J., Reed, L.K., 2016. Genetic and sex-specific transgenerational effects of a high fat diet in *Drosophila melanogaster*. PLoS One 11, e0160857.

17. Drijfhout, F.P., Kather, R., Martin, S.J., 2009. The role of cuticular hydrocarbons in insects. Behavioral and chemical ecology 91–114.

18. Edgerly, J.S., Mcfarland, M., Morgan, P., Livdahl, T., 1998. A seasonal shift in egg-laying behaviour in response to cues of future competition in a treehole mosquito. J. Anim. Ecol. 67, 805–818.

19. Farnesi, L.C., Vargas, H.C.M., Valle, D., Rezende, G.L., 2017. Darker eggs of mosquitoes resist more to dry conditions: Melanin enhances serosal cuticle contribution in egg resistance to desiccation in *Aedes, Anopheles* and *Culex* vectors. PLoS Negl. Trop. Dis. 11, e0006063.

20. Faull, K.J., Webb, C., Williams, C.R., 2016. Desiccation survival time for eggs of a widespread and invasive Australian mosquito species, *Aedes (Finlaya) notoscriptus (Skuse)*. J. Vector Ecol. 41, 55–62.

21. Fischer, K., Brakefield, P.M., Zwaan, B.J., 2003. Plasticity in butterfly egg size: Why larger offspring at lower temperatures? Ecology 84, 3138–3147.

22. Fox, C.W., Czesak, M.E., Mousseau, T.A., Roff, D.A., 1999. The evolutionary genetics of an adaptive maternal effect: Egg size plasticity in a seed beetle. Evolution 53, 552–560.

23. Gefen, E., Marlon, A.J., Gibbs, A.G., 2006. Selection for desiccation resistance in adult *Drosophila melanogaster* affects larval development and metabolite accumulation. J. Exp. Biol. 209, 3293–3300.

24. Hans, K.R., LeBouthillier, R., VanLaerhoven, S.L., 2019. Effect of temperature on oviposition behavior and egg load of blow flies (Diptera: Calliphoridae). J. Med. Entomol. 56, 441–447.

25. Helinski, M.E.H., Harrington, L.C., 2011. Male mating history and body size influence female fecundity and longevity of the dengue vector *Aedes aegypti*. J. Med. Entomol. 48, 202–211.

26. Hilker, M., Salem, H., Fatouros, N.E., 2023. Adaptive plasticity of insect eggs in response to environmental challenges. Annu. Rev. Entomol. 68, 451–469.

27. Hoffmann, A.A., Hallas, R., Sinclair, C., Partridge, L., 2001. Rapid loss of stress resistance in *Drosophila melanogaster* under adaptation to laboratory culture. Evolution 55, 436–438.

28. Holmes, C.J., Benoit, J.B., 2019. Biological adaptations associated with dehydration in mosquitoes. Insects 10, 375.

29. Holmes, C.J., Brown, E.S., Sharma, D., Warden, M., Pathak, A., Payton, B., Nguyen, Q., Spangler, A., Sivakumar, J., Hendershot, J.M., Benoit, J.B., 2023. Dehydration alters transcript levels in the mosquito midgut, likely facilitating rapid rehydration following a bloodmeal. Insects 14. 10.3390/insects14030274

30. Holmes, C.J., Chakraborty, S., Ajayi, O.M., Uhran, M.R., Frigard, R., Stacey, C.L., Susanto, E.E., Chen, S.-C., Rasgon, J.L., DeGennaro, M., Xiao, Y., Benoit, J.B., 2025. Multiple blood feeding bouts in mosquitoes allow for prolonged survival and are predicted to increase viral transmission during dry periods. iScience 28, 111760.

31. Hopper, K.R., 1999. Risk-spreading and bet-hedging in insect population biology. Annu. Rev. Entomol. 44, 535–560.

32. J. Marshall, D., Uller, T., 2007. When is a maternal effect adaptive? Oikos 116, 1957–1963.

33. Juliano, S.A., O’Meara, G.F., Morrill, J.R., Cutwa, M.M., 2002. Desiccation and thermal tolerance of eggs and the coexistence of competing mosquitoes. Oecologia 130, 458–469.

34. Kearney, M., Porter, W.P., Williams, C., Ritchie, S., Hoffmann, A.A., 2009. Integrating biophysical models and evolutionary theory to predict climatic impacts on species’ ranges: the dengue mosquito *Aedes aegypti* in Australia. Funct. Ecol. 23, 528–538.

35. Lacour, G., Vernichon, F., Cadilhac, N., Boyer, S., Lagneau, C., Hance, T., 2014. When mothers anticipate: effects of the prediapause stage on embryo development time and of maternal photoperiod on eggs of a temperate and a tropical strains of *Aedes albopictus* (Diptera: Culicidae). J. Insect Physiol. 71, 87–96.

36. Ledón-Rettig, C.C., 2023. A transcriptomic investigation of heat-induced transgenerational plasticity in beetles. Biol. J. Linn. Soc. Lond. 138, 318–327.

37. Le Hesran, S., Groot, T., Knapp, M., Bukovinszky, T., Forestier, T., Dicke, M., 2019. Phenotypic variation in egg survival in the predatory mite *Phytoseiulus persimilis* under dry conditions. Biol. Control 130, 88–94.

38. Le Hesran, S., Groot, T., Knapp, M., Bukovinszky, T., Nugroho, J.E., Beretta, G., Dicke, M., 2020. Maternal effect determines drought resistance of eggs in the predatory mite *Phytoseiulus persimilis*. Oecologia 192, 29–41.

39. Le Roux, L., Meunier, J., Villalta, I., 2024. Heat waves during egg development alter maternal care and offspring quality in the European earwig. J. Therm. Biol. 125, 104006.

40. Li, A., Benoit, J.B., Lopez-Martinez, G., Elnitsky, M.A., Lee, R.E., Jr, Denlinger, D.L., 2009. Distinct contractile and cytoskeletal protein patterns in the Antarctic midge are elicited by desiccation and rehydration. Proteomics 9, 2788–2798.

41. Lopez-Martinez, G., Benoit, J.B., Rinehart, J.P., Elnitsky, M.A., Lee, R.E., Jr, Denlinger, D.L., 2009. Dehydration, rehydration, and overhydration alter patterns of gene expression in the Antarctic midge, *Belgica antarctica*. J. Comp. Physiol. B 179, 481–491.

42. Loveridge, J.P., Bursell, E., 1975. Studies on the water relations of adult locusts (orthoptera, acrididae). I. Respiration and the production of metabolic water. Bull. Entomol. Res. 65, 13– 20.

43. Macagno, A.L.M., Zattara, E.E., Ezeakudo, O., Moczek, A.P., Ledón-Rettig, C.C., 2018. Adaptive maternal behavioral plasticity and developmental programming mitigate the transgenerational effects of temperature in dung beetles. Oikos 127, 1319–1329.

44. Mamai, W., Mouline, K., Blais, C., Larvor, V., Dabiré, K.R., Ouedraogo, G.A., Simard, F., Renault, D., 2014. Metabolomic and ecdysteroid variations in *Anopheles gambiae* s.l. mosquitoes exposed to the stressful conditions of the dry season in Burkina Faso, West Africa. Physiol. Biochem. Zool. 87, 486–497.

45. Mogi, M., Tuno, N., 2014. Impact of climate change on the distribution of *Aedes albopictus* (Diptera: Culicidae) in northern Japan: retrospective analyses. J. Med. Entomol. 51, 572– 579.

46. Mousseau, T.A., Dingle, H., 1991. Maternal effects in insect life histories. Annu. Rev. Entomol. 36, 511–534.

47. Mousseau, T.A., Fox, C.W., 1998. The adaptive significance of maternal effects. Trends Ecol. Evol. 13, 403–407.

48. Nussey, D.H., Wilson, A.J., Brommer, J.E., 2007. The evolutionary ecology of individual phenotypic plasticity in wild populations. J. Evol. Biol. 20, 831–844.

49. Pigliucci, M., 2005. Evolution of phenotypic plasticity: where are we going now? Trends Ecol. Evol. 20, 481–486.

50. Potter, K.A., Woods, H.A., 2012. No evidence for the evolution of thermal or desiccation tolerance of eggs among populations of *Manduca sexta*: Egg size and abiotic stress. Funct. Ecol. 26, 112–122.

51. Prasad, A., Sreedharan, S., Bakthavachalu, B., Laxman, S., 2023. Eggs of the mosquito *Aedes aegypti* survive desiccation by rewiring their polyamine and lipid metabolism. PLoS Biol. 21, e3002342.

52. Reinhold, J.M., Lazzari, C.R., Lahondère, C., 2018. Effects of the Environmental Temperature on *Aedes aegypti* and *Aedes albopictus* Mosquitoes: A Review. Insects 9. 10.3390/insects9040158

53. Rezende, G.L., Martins, A.J., Gentile, C., Farnesi, L.C., Pelajo-Machado, M., Peixoto, A.A., Valle, D., 2008. Embryonic desiccation resistance in Aedes aegypti: presumptive role of the chitinized serosal cuticle. BMC Dev. Biol. 8, 82.

54. Rosendale, A.J., Romick-Rosendale, L.E., Watanabe, M., Dunlevy, M.E., Benoit, J.B., 2016. Mechanistic underpinnings of dehydration stress in the American dog tick revealed through RNA-Seq and metabolomics. J. Exp. Biol. 219, 1808–1819.

55. Rose, N.H., Sylla, M., Badolo, A., Lutomiah, J., Ayala, D., Aribodor, O.B., Ibe, N., Akorli, J., Otoo, S., Mutebi, J.-P., Kriete, A.L., Ewing, E.G., Sang, R., Gloria-Soria, A., Powell, J.R., Baker, R.E., White, B.J., Crawford, J.E., McBride, C.S., 2020. Climate and urbanization drive mosquito preference for humans. Curr. Biol. 30, 3570–3579.e6.

56. Rossiter, M.C., 1991. Environmentally-based maternal effects: a hidden force in insect population dynamics? Oecologia 87, 288–294.

57. Saifur, R.G.M., Dieng, H., Hassan, A.A., Satho, T., Miake, F., Boots, M., Salmah, R.C., Abubakar, S., 2010. The effects of moisture on ovipositional responses and larval eclosion of *Aedes albopictus*. J. Am. Mosq. Control Assoc. 26, 373–380.

58. Schausberger, P., 1998. The influence of relative humidity on egg hatch in *Euseius filandicus, Typhlodromus pyri and Kampimodromus aberrans (Acari, Phytoseiidae)*. J. Appl. Entomol. 122, 497–500.

59. Shaman, J., Day, J.F., 2007. Reproductive phase locking of mosquito populations in response to rainfall frequency. PLoS One 2, e331.

60. Simons, A.M., 2011. Modes of response to environmental change and the elusive empirical evidence for bet hedging. Proc. Biol. Sci. 278, 1601–1609.

61. Sinclair, B.J., Gibbs, A.G., Roberts, S.P., 2007. Gene transcription during exposure to, and recovery from, cold and desiccation stress in *Drosophila melanogaster*. Insect Mol. Biol. 16, 435–443.

62. Sirot, L.K., Hardstone, M.C., Helinski, M.E.H., Ribeiro, J.M.C., Kimura, M., Deewatthanawong, P., Wolfner, M.F., Harrington, L.C., 2011. Towards a semen proteome of the dengue vector mosquito: protein identification and potential functions. PLoS Negl. Trop. Dis. 5, e989.

63. Sota, T., Mogi, M., 1992. Interspecific variation in desiccation survival time of *Aedes (Stegomyia)* mosquito eggs is correlated with habitat and egg size. Oecologia 90, 353–358.

64. Sturiale, S.L., Bailey, N.W., 2023. Within-generation and transgenerational social plasticity interact during rapid adaptive evolution. Evolution 77, 409–421.

65. Telonis-Scott, M., Guthridge, K.M., Hoffmann, A.A., 2006. A new set of laboratory-selected *Drosophila melanogaster* lines for the analysis of desiccation resistance: response to selection, physiology and correlated responses. J. Exp. Biol. 209, 1837–1847.

66. Vargas, H.C.M., Farnesi, L.C., Martins, A.J., Valle, D., Rezende, G.L., 2014. Serosal cuticle formation and distinct degrees of desiccation resistance in embryos of the mosquito vectors *Aedes aegypti*, *Anopheles aquasalis* and *Culex quinquefasciatus*. J. Insect Physiol. 62, 54– 60.

67. Venkataraman, K., Shai, N., Lakhiani, P., Zylka, S., Zhao, J., Herre, M., Zeng, J., Neal, L.A., Molina, H., Zhao, L., Vosshall, L.B., 2023. Two novel, tightly linked, and rapidly evolving genes underlie *Aedes aegypti* mosquito reproductive resilience during drought. Elife 12, e80489.

68. Walzer, A., Schausberger, P., 2015. Food stress causes sex-specific maternal effects in mites. J. Exp. Biol. 218, 2603–2609.

69. Whitman, D., Agrawal, A., 2009. What is Phenotypic Plasticity and Why is it Important?, in: Phenotypic Plasticity of Insects. Science Publishers.

70. Woestmann, L., Saastamoinen, M., 2016. The importance of trans-generational effects in Lepidoptera. Curr. Zool. 62, 489–499.

71. Yanchula, K.Z., Alto, B.W., 2021. Paternal and maternal effects in a mosquito: A bridge for life history transition. J. Insect Physiol. 131, 104243.

72. Yoder, J.A., Benoit, J.B., Opaluch, A.M., 2004. Water relations in eggs of the lone star tick, *Amblyomma americanum*, with experimental work on the capacity for water vapor absorption. Exp. Appl. Acarol. 33, 235–242.

73. Yoder, J.A., Tank, J.L., Rellinger, E.J., 2006. Evidence of a maternal effect that protects against water stress in larvae of the American dog tick, *Dermacentor variabilis* (Acari: Ixodidae). J. Insect Physiol. 52, 1034–1042.

74. Zanchi, C., Troussard, J.-P., Martinaud, G., Moreau, J., Moret, Y., 2011. Differential expression and costs between maternally and paternally derived immune priming for offspring in an insect: Male vs. female immune priming for offspring. J. Anim. Ecol. 80, 1174–1183.

75. Zhou, J.-C., Liu, Q.-Q., Han, Y.-X., Dong, H., 2018. High temperature tolerance and thermal- adaptability plasticity of Asian corn borer (*Ostrinia furnacalis* Guenée) after a single extreme heat wave at the egg stage. J. Asia. Pac. Entomol. 21, 1040–1047.

76. Zirbel, K.E., Alto, B.W., 2018. Maternal and paternal nutrition in a mosquito influences offspring life histories but not infection with an arbovirus. Ecosphere 9, e02469.

