## Supplementary figures and images for "Parental exposure to wet and dry conditions shapes the viability and thermotolerance of eggs in *Aedes aegypti*"

### Supplemental Figure 1

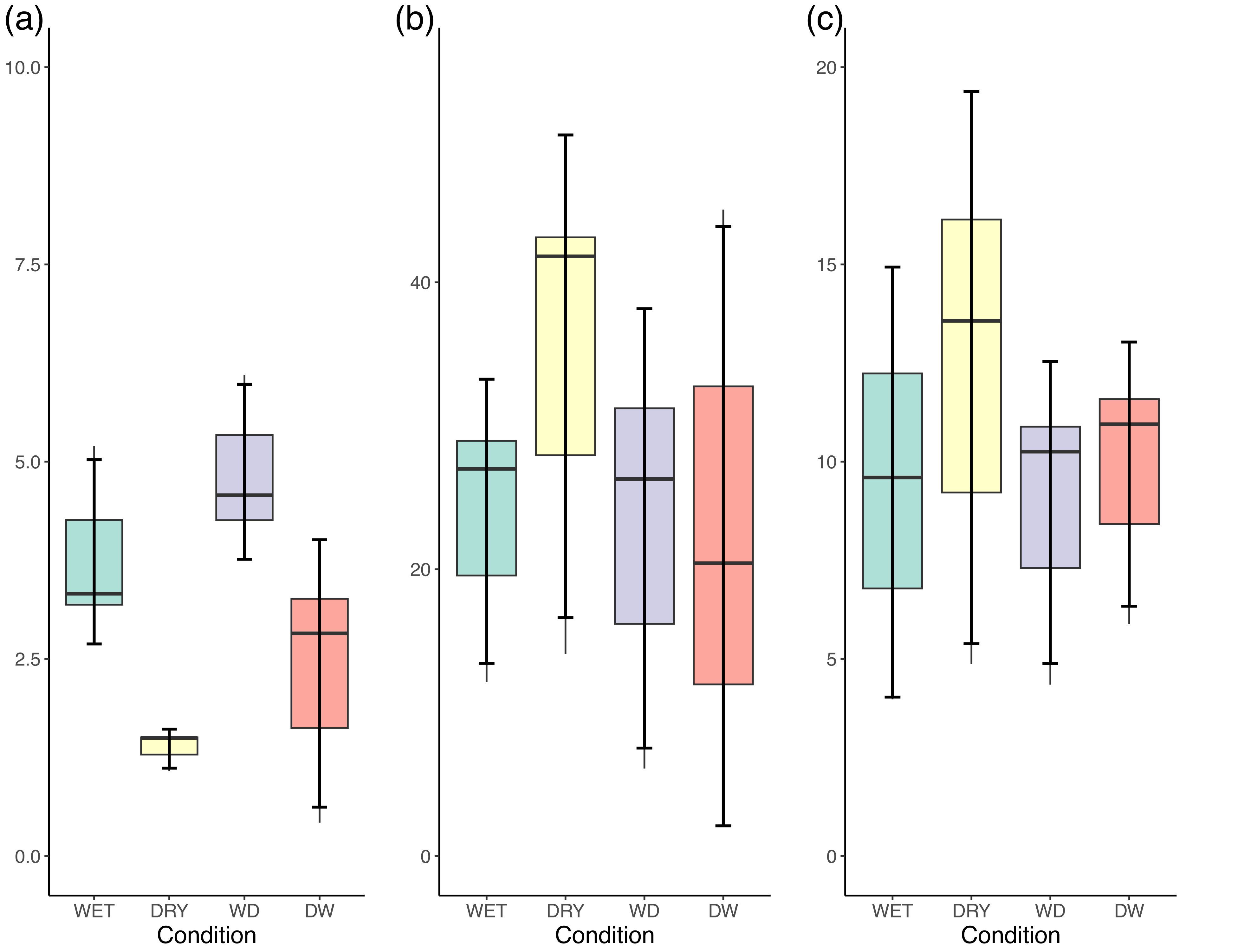
